# A quantitative three-dimensional bone mineralisation assay using human dental pulp stem cell spheroids

**DOI:** 10.64898/2026.10.27.762162

**Authors:** Mohammed Alshadidi, Elad Katz, Ian Ellis, Sarah Jones, Mohammad Islam

## Abstract

Matric mineralisation in three-dimensional (3D) human cell culture is still assessed predominantly by terminal histochemical staining. Alizarin Red S and Von Kossa are destructive, semi-quantitative, and require fixation and dye extraction before a number can be obtained, which precludes longitudinal measurement and makes miniaturised multiwell formats impractical. We describe a scaffold-free assay that renders 3D human bone mineralisation as a continuous, well-level quantity. Primary human dental pulp stem cells (DPSCs) were dispensed by automated liquid handling into 384-well plates, where they formed a single, centrally positioned spheroid per well. Eleven plate geometry and surface-coating combinations were compared, and cell-repellent coating rather than well geometry alone was identified as the determinant of single-aggregate formation. Mineral deposition was measured non-destructively by adding the tetracycline-derived near-infrared probe BoneTag680 directly to the existing culture volume and reading the plate from below, with no spheroid transfer, embedding, sectioning or dye extraction at any point. Spheroids maintained in 10% fetal calf serum (FCS) with osteogenic cocktail were separated from a 1% FCS low-mineralisation condition with an assay window of 13.74 ± 3.29-fold and a Z′ factor of 0.627 ± 0.119 across four independent plates, with a well-level coefficient of variation of 10.88% compared with 55.81% in a commercial alternative. A DNAcontent measurement acquired from the same wells after near-infrared acquisition showed no increase in cell number under osteogenic conditions, establishing that the mineral signal is not biomass-driven and providing an internal counter-screen against proliferative and cytotoxic artefacts. All data derive from a single donor; the assay is therefore characterised for technical and within-donor reproducibility, and is presented as a defined quantitative measurement intended as a foundation for bone-anabolic compound

## Introduction

Bone anabolic discovery is limited less by the availability of candidate compounds than by the assays used to evaluate them. Osteogenic differentiation and matrix mineralisation are still assessed predominantly in two-dimensional (2D) monolayers and scored by terminal histochemistry. Both conventions impose real constraints. Planar culture removes the cell-cell and cell-matrix geometry that governs stem cell lineage commitment and matrix organisation,^1,2^ and terminal staining converts a dynamic process into a single destructive measurement, requiring dye extraction or image segmentation before a number can be obtained. Mineralisation - the phenotype of interest - is therefore rarely measured as a continuous, well-level quantity in a human three-dimensional (3D) context.

Human 3D models have been developed to restore that missing architecture and to create systems in which cellular organisation and function can be interrogated experimentally.^3^ Dental pulp offers a practical and scalable human cell source for mineralised tissue models. Dental pulp stem cells (DPSCs) were identified as a clonogenic, rapidly proliferating population competent to form mineralised tissue,^4^ with demonstrated self-renewal and multilineage potential^5^ and a mesenchymal stem cell-associated surface phenotype.^6^ Unlike bone marrow-derived stem cells, DPSCs are recoverable non-invasively from routinely extracted third molars that would otherwise be discarded and are amenable to banking and cryopreservation.^7^ Scaffold-free DPSC and mesenchymal stem cell spheroids undergo odontogenic and osteogenic differentiation and support bone regeneration.^8–10^ These studies establish the biology but do not solve the assay problem, having been designed to demonstrate differentiation or regenerative capacity rather than to generate a quantitative signal at well level.

Two barriers stand between 3D culture and a usable assay. First, spheroid behaviour is sensitive to methodological variables. Systematic analysis of more than 3,000 published spheroid experiments identified pervasive heterogeneity in aggregation method, culture geometry and seeding density, and showed that such variation alters measured spheroid metrics;^11^ workflows for automated spheroid handling identify morphological and positional consistency as a precondition for reproducibility.^12^ In a multiwell plate, an aggregate varying in size, integrity or position introduces variance directly into the optical readout. Second, the endpoint must become non-destructive and continuous. Tetracycline-derived bone-targeting probes render mineral deposition as a measurable optical signal rather than a histological observation,^13,14^ and 3D models have been adapted to quantitative multiwell workflows once culture generation, handling and detection are sufficiently controlled.^15^ Design considerations for screen-compatible osteogenesis models have been set out specifically for this purpose.^16^ Assay miniaturisation becomes meaningful only once phenotypic consistency and signal separation are established.

Here we describe a scaffold-free 3D human DPSC mineralisation assay in 384-well format with a non-destructive near-infrared readout. We established osteogenic responsiveness in 2D, compared eleven plate geometry and coating combinations to identify conditions producing a single, centrally positioned spheroid per well, and defined a pair of culture conditions bracketing the measurable signal range. Mineralisation was quantified by direct in-well labelling and plate-reader fluorescence, with whole-plate imaging used to assess spatial uniformity and a DNA-content measurement from the same wells to distinguish mineral signal from biomass.

## Methods

### Human tissue collection

DPSCs were isolated from freshly extracted human third molars that would otherwise have been discarded following routine clinical procedures. Samples were collected with informed consent under Tayside Biorepository delegated ethics (Research Tissue Collection, approval no. TR000620). Teeth were obtained from the Oral Surgery Clinic at Dundee Dental Hospital and School and the Broxden Dental Outreach Centre. Inclusion required partially or fully erupted, unrestored, non-carious teeth free from periapical or periodontal disease. Teeth were transferred under sterile conditions in phosphate-buffered saline (PBS) with penicillin–streptomycin and amphotericin B and processed immediately. **All data reported in this study were generated from a single DPSC donor**. Additional donors were isolated during the wider project but used in separate experiments under different conditions and are not reported here.

### Isolation, culture and cryopreservation

Pulp tissue was aseptically harvested from sectioned teeth, cut into explants and transferred to tissue-culture-treated vessels for adherent outgrowth.^17,18^ Cultures were maintained in α-MEM with 10% fetal calf serum (FCS), 1% penicillin–streptomycin and 2.5 µg/mL amphotericin B at 37°C in 5% CO_2_, with medium changed every 2–3 days, and passaged at 80–90% confluence using trypsin-EGTA. Mycoplasma contamination was routinely assessed by a standard colorimetric assay. Aliquots were cryopreserved in standard freezing medium and stored in liquid nitrogen.^7^ All experiments used cells between passages 3 and 6 to limit passage-associated phenotypic drift.^19^ Reagents, antibodies, instruments and software are listed in Table 1.

### Immunophenotyping and colony-forming assay

Multicolour flow cytometry used mesenchymal stem cell-associated and lineage-negative markers.^6,18^ Cells were detached, resuspended in PBS with 1% FCS and stained at 5 × 10^5^ cells per sample in 100 µL on ice, protected from light. Viability was assessed by DAPI exclusion. Compensation used BD CompBeads with unstained and single-stained controls. Data were acquired on a BD LSRFortessa configured with 488, 561 and 640 nm lasers (FITC and PerCP-Cy5.5 from 488 nm; PE and PE-Cy7 from 561 nm; APC and APC/Fire 750 from 640 nm) and analysed in FlowJo v10.8.1 using sequential gating for population, viability and singlets. For colony-forming unit-fibroblast assay, cells were plated at 1,000 per 60 mm dish for 7 days, fixed, stained with 1% toluidine blue, and colonies of ≥25 cells counted.^20^ Colony-forming efficiency was colonies formed divided by cells seeded, as a percentage.^4^

### Culture conditions

**2D**. DPSCs were seeded in α-MEM with 10% FCS. After attachment, cultures were assigned to 10% FCS, or to 10% FCS + OC (α-MEM, 10% FCS, and an osteogenic cocktail comprising 100 nM dexamethasone, 50 µM L-ascorbic acid and 10 mM β-glycerophosphate).^21,22^ Both conditions contained 10% FCS and were maintained up to 21 days with medium replaced every 2–3 days.

**3D**. Spheroids were seeded on day 0 in α-MEM with 1% FCS. On day 1, medium was exchanged to define the two assay conditions: 1% FCS, the low-mineralisation anchor, and 10% FCS + OC, the maximal-mineralisation anchor. Spheroids were then maintained without further medium exchange until endpoint, minimising handling-associated variability.

#### Derivation of the assay anchors

Both conditions were established empirically. The low anchor was required to give minimal mineral signal while maintaining spheroid integrity and cell number across the 5-7 day assay period. Initial experiments with 10% FCS in both arms produced variable signal in the unsupplemented arm and a narrow assay window; complete serum withdrawal (0% FCS) compromised spheroid integrity and caused disaggregation; 1% FCS without osteogenic cocktail maintained aggregate integrity and cell number while suppressing mineral signal. The high anchor was required to give maximal mineral signal without an increase in cell number, which 10% FCS + OC met. These conditions are used throughout as anchors bracketing the measurable signal range, and do not constitute a serum-matched comparison isolating the effect of the osteogenic cocktail.

### Spheroid generation, plate comparison and automated handling

DPSCs were resuspended as single cells and dispensed at 500–8,000 cells per well in 40 µL. Optimisation examined the full range; 2,000 cells per well was selected as a compromise between conserving primary human material and maintaining an aggregate robust enough for repeated optical measurement. Plates were incubated under standard conditions to permit self-assembly, with morphology monitored immediately after seeding and at a 24-h quality-control checkpoint. Aggregation was compared across the plate formats in Table 1, coated and uncoated where applicable; coating used BIOFLOAT FLEX, 50 µL per well for 5 min, removed and air-dried 2 h under sterile conditions. This comparison was a qualitative screen: the operational expectation at the checkpoint was an intact, centrally localised aggregate, and quantitative acceptance criteria based on circularity, projected area or diameter were not defined prospectively. No retrospective morphometric analysis was performed. 384-well V-bottom plates were excluded because well-wall optical artefacts prevented reliable bright-field morphological assessment. Automated workflows used a FluidX XPP-721 (96-tip head, manual tip configuration) and an INTEGRA Assist Plus (VIAFLO 125 µL 8-channel head), with protocols built in NMS and VIALAB respectively, for plate coating, cell dispensing and medium exchange.

### Immunofluorescence

**2D**. Cultures were fixed at days 7, 14 and 21 in 4% paraformaldehyde, permeabilised in 0.2% Triton X-100 for 5 min and blocked 1 h in 5% normal goat serum in 0.1% Tween-PBS. Anti-COL1α1 (1:200) was applied overnight, followed by anti-rabbit Alexa Fluor 488 (1:500) and DAPI. Imaging used an Olympus IX70 under matched acquisition and display settings.^23^

**3D**. Spheroids were processed in-plate: fixation in 4% paraformaldehyde for 20 min at room temperature, permeabilisation in 40 µL 0.5% Triton X-100 in PBS for 30 min, blocking 1 h in 40 µL 2% BSA with 0.1% Triton X-100. Anti-COL1α1 or anti-osteocalcin (1:200) was applied overnight, followed by secondary antibody and DAPI. Wash steps minimised spheroid displacement. Imaging used a CV7000 high-content system under matched acquisition settings.^9^ Images are representative; no intensity quantification was performed.

### Alizarin Red S and BoneTag680 labelling

Alizarin Red S was applied to 2D cultures at days 7, 14 and 21 after fixation in 4% paraformaldehyde, for 20 min at room temperature, with excess dye removed by serial washes in distilled water, and used as a qualitative reference for mineral deposition.^4,22^ For 2D BoneTag680 labelling, IRDye 680RD BoneTag^13^ was added directly to culture medium at 1:1,000 final dilution 24 h before each endpoint; cultures were washed, fixed in 20% neutral buffered formalin and scanned on an Odyssey CLx, with images analysed in Image Studio under matched settings. For 3D, 10 µL BoneTag680 working solution was added on day 4 to the existing 40 µL culture volume, giving 50 µL at 1:500 final dilution; on day 5 spheroids were washed in PBS and fixed in 20% neutral buffered formalin for 20 min before acquisition.

### Fluorescence acquisition, whole-plate imaging and DNA content

BoneTag680 fluorescence was measured on a PHERAstar FS using fluorescence-intensity acquisition with 660–20 nm excitation and 705–20 nm emission filters and bottom optics; gain was optimised against a high-signal 10% FCS + OC well with automatic adjustment to 35% target. Whole-plate near-infrared images were acquired on an Odyssey CLx in the 700 nm channel at 169 µm resolution with 4.0 mm focus offset, and used to assess spatial signal distribution; PHERAstar measurements were the primary quantitative endpoint. DNA content was measured using CyQUANT NF adapted for 3D spheroids: following BoneTag680 acquisition, wells were equilibrated in HBSS, dye prepared at 2× in 1× HBSS was added, and plates incubated 30 min at room temperature in the dark before measurement. Blank wells containing buffer and dye without spheroids were included on every plate and net values calculated after blank subtraction.

### Mechanical stimulation

Plates were placed on a Luckham R100 orbital shaker at approximately 55 rpm for 4 h per day following addition of the osteogenic cocktail. After 5 days, spheroids were fixed and stained for osterix (1:200) and sclerostin (1:200)^24^ with DAPI counterstain and imaged on a CV7000 under matched settings. The hydrodynamic environment was not characterised.

### Statistical analysis

Analyses were performed in GraphPad Prism 10. All data derive from a single DPSC donor. “Independent experiment” denotes a separate plate seeded on a separate day from an independently thawed cell stock; the number of wells and independent experiments is stated in each figure legend. Because all data derive from one donor, replicate wells are technical replicates and all reported statistics describe technical and within-donor variability; no inference about biological variability between donors is made. Assay quality was assessed by coefficient of variation and Z′ factor, calculated as Z′ = 1 − [3(σ~10% FCS + OC~ + σ~1% FCS_)/|µ_10% FCS + OC~ − µ~1% FCS~|], across four independent runs. Variances between plate formats were compared by F-test. Significance was defined as p < 0.05.

### Experiment

#### An automated 384-well workflow requiring no spheroid transfer

The assay was designed around a single constraint: every step from seeding to readout had to be executable without removing the spheroid from its well. Aggregate transfer is a dominant source of loss and positional variance in 3D multiwell workflows,^12^ and each transfer compounds well-to-well variability in the final optical measurement. DPSCs are dispensed into coated 384-well plates by automated liquid handling and allowed to self-assemble. Wells are assigned to 1% FCS or 10% FCS + OC at 24 h. BoneTag680 is added directly to the existing culture volume and mineral-associated signal acquired by bottom-read plate-reader fluorescence, with whole-plate near-infrared imaging providing spatial context from the same physical plate (Figure 1).

**Figure 1.**
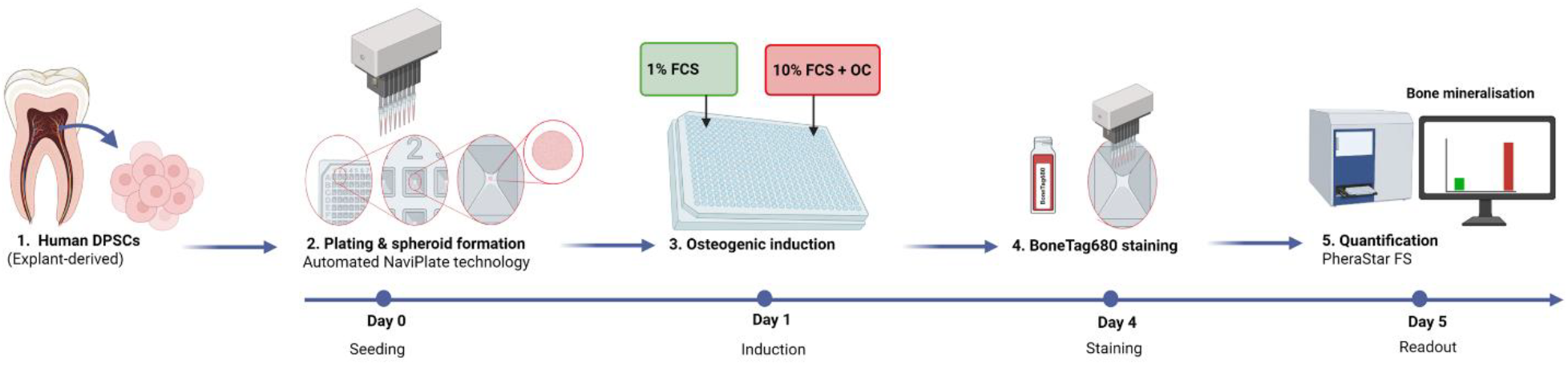
Automated 384-well workflow for quantitative mineralisation in human DPSC spheroids. (A) Overview. (B) Human DPSCs are dispensed into 384-well plates by automated liquid handling to generate a single spheroid per well; inset shows well geometry and aggregate position. (C) After spheroid establishment, wells are assigned to 1% FCS or 10% FCS + OC. (D, E) BoneTag680 is added directly to the culture and mineral-associated near-infrared signal quantified by plate reader, with whole-plate imaging acquired from the same plate. Created with BioRender.

#### Explant-derived DPSCs display clonogenic capacity and an MSC-associated immunophenotype

DPSCs were derived by explant outgrowth, yielding adherent fibroblast-like populations that migrated from pulp tissue within 7-10 days (Figure 2A). Clonogenic capacity was confirmed by colony-forming unit-fibroblast assay, with discrete colonies visible after toluidine blue staining (Figure 2B) and a colony-forming efficiency of approximately 17% (Figure 2C). Flow cytometry established an MSC-associated surface profile: 98% of events were CD90^+^ and lineage-negative, 99.9% CD13^+^CD73^+^, and 93.9% CD105^+^CD29^+^ (Figure 2D), meeting accepted minimal criteria.^6^ These data derive from a single donor and establish the identity of the population used for assay development; they are not intended to describe DPSC populations generally.

**Figure 2.**
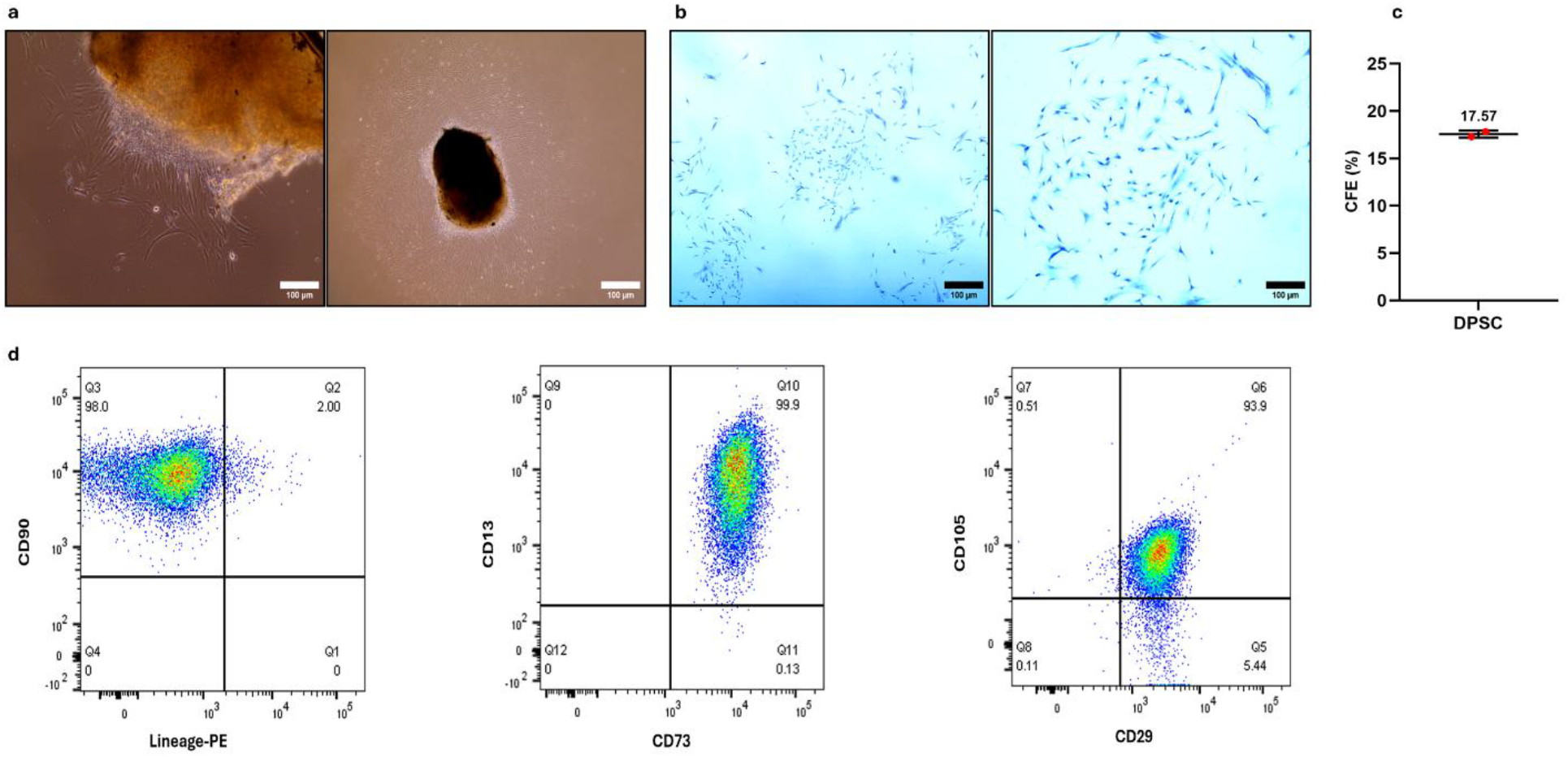
Explant-derived human DPSCs exhibit clonogenic capacity and an MSC-like immunophenotype. All data are from a single DPSC donor. (A) DPSC outgrowth at day 7 from dental pulp explants. (B) CFU-F assay, toluidine blue-stained colonies at clonal density. (C) Colony-forming efficiency (%). (D) Flow cytometry dot plots for CD90, CD13, CD73, CD105 and CD29, with a lineage-negative gate (CD34/CD45/HLA-DR). Scale bars, 100 μm (A, B).

#### Near-infrared labelling reports mineralisation in 2D culture

Before developing the 3D format, we established that the probe reports mineralisation in monolayer culture, where Alizarin Red S provides a reference standard. Both 2D conditions contained 10% FCS, differing only in osteogenic cocktail. COL1α1 was detected in both conditions at all timepoints, confirming that collagen expression alone does not discriminate osteogenic induction in this system (Figure 3A). Mineralisation-associated readouts separated the conditions: Alizarin Red S was low in both groups at day 7, increased substantially in 10% FCS + OC by day 14 and remained elevated at day 21, while 10% FCS cultures stained weakly throughout (Figure 3B). BoneTag680 followed the same trajectory (Figure 3C), reaching 147%, 5,871% and 7,745% of the 10% FCS condition at days 7, 14 and 21 (Figure 3D). Because osteogenic supplementation alone produced clear separation at equal serum concentration, no serum reduction was applied in the monolayer system.

**Figure 3.**
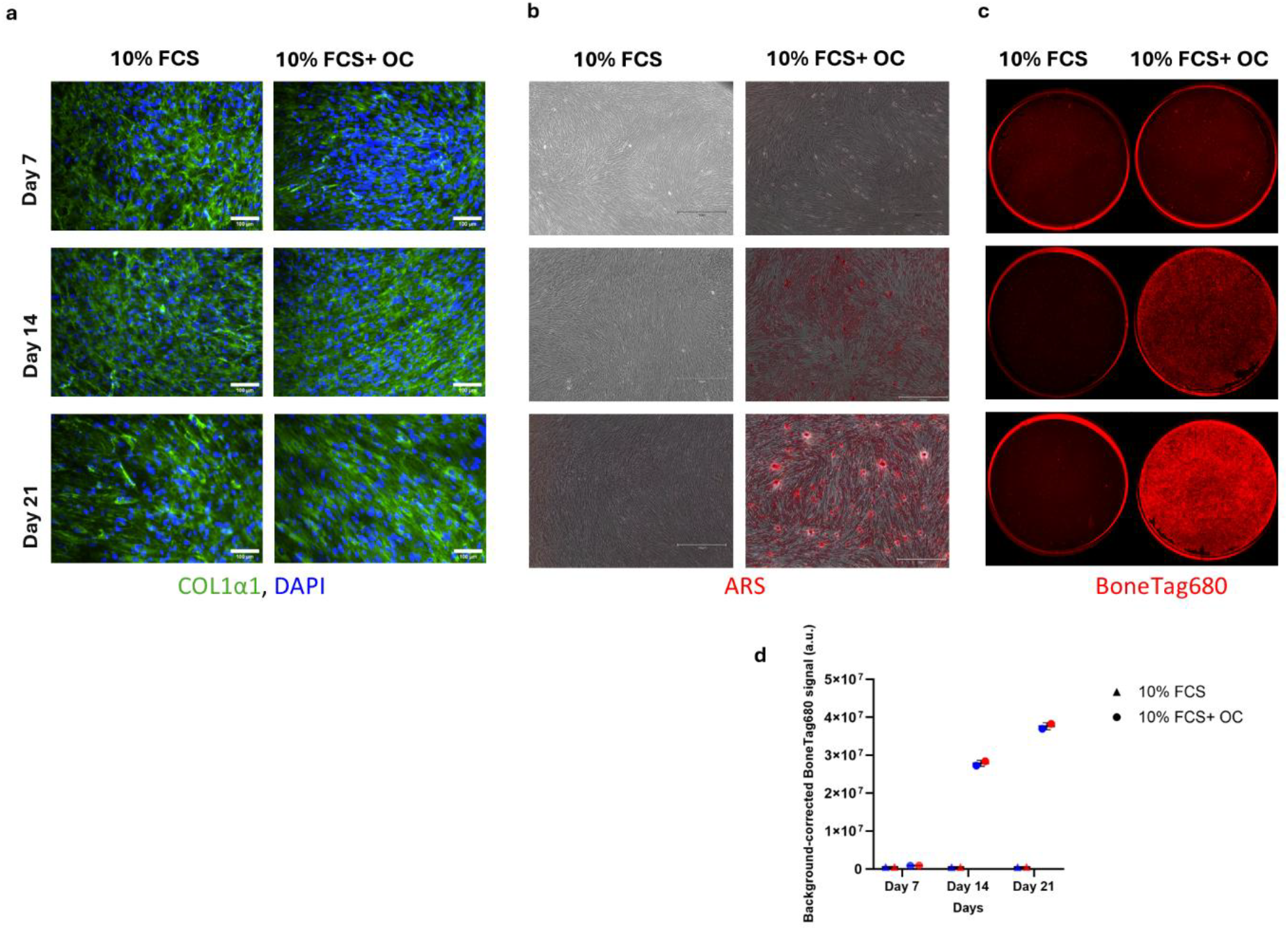
Time-course of osteogenic differentiation in 2D DPSC culture. DPSCs under 10% FCS or 10% FCS+ OC at days 7, 14 and 21. (A) COL1α1 immunofluorescence (green) with DAPI (blue). (B) Alizarin Red S. (C) Odyssey CLx scans of BoneTag680-labelled cultures. (D) BoneTag680 signal expressed as a percentage of the 10% FCS condition; dashed line, 10% FCS level (100%). n = 2 independent experiments, single donor. Scale bars, 750 µm.

#### Surface coating and geometry determines single-spheroid formation

Eleven plate geometry and coating combinations were compared under matched seeding density, dispense method and incubation conditions (Figure 4A–K). 96-well U-bottom plates gave dispersed adherent cells with no aggregate in either coating condition (Figure 4A,B); 96-well V-bottom plates produced partial aggregation with heterogeneous morphology (Figure 4C,D). 384-well V-bottom plates were excluded, as optical artefacts prevented reliable morphological assessment, and no conclusion about that format should be drawn from this study (Figure 4E,F). Microcavity plates distributed cells across multiple subwells, generating many micro-aggregates rather than one (Figure 4G,H) - appropriate where aggregate populations are wanted, but incompatible with a well-level optical readout. Two formats produced a single consolidated aggregate: 384-well U-bottom BioFloat (Figure 4I) and coated NaviPlate (Figure 4K). Uncoated NaviPlate wells gave fragmented or multiple aggregates (Figure 4J), identifying cell-repellent surface coating rather than well geometry alone as the determining variable - the clearest transferable observation from this comparison. Both coated formats were carried forward for quantitative assessment.

**Figure 4.**
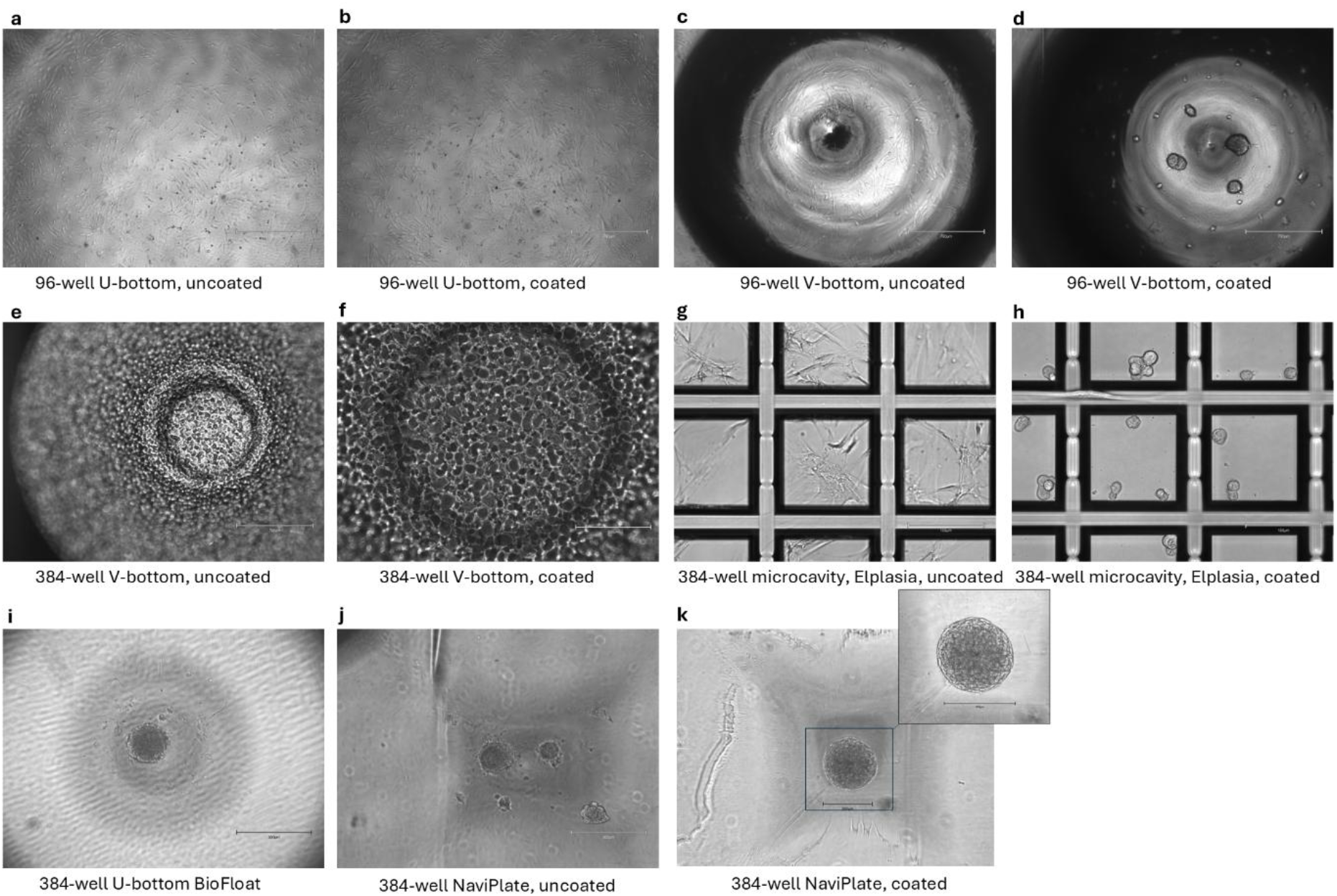
Qualitative comparison of plate formats for single-spheroid formation. Representative bright-field images of DPSC aggregation 24 h after seeding under matched seeding density, dispense method and incubation conditions. (A–D) 96-well U-bottom and V-bottom, uncoated and coated. (E, F) 384-well Vbottom, uncoated and coated. (G, H) 384-well microcavity, uncoated and coated. (I) 384-well U-bottom BioFloat. (J, K) 384-well NaviPlate, uncoated and coated. (L) Developed NaviPlate spheroid after continued culture. This comparison was performed as a qualitative screen; no prospective quantitative acceptance criteria were defined and no morphometric analysis was performed. Scale bars, 300 μm (384-well plates) and 750 µm (96-well plates).

#### The selected format gives a more consistent mineralisation signal

Both candidate formats supported early matrix production, with COL1α1 detectable in 3D aggregates within 24 h at levels comparable to monolayers (Figure 5A). Mineral deposition was detectable within 5 days of osteogenic cocktail addition. BoneTag680 signal normalised to the 1% FCS condition was 1819.6 ± 198.0% in NaviPlate and 2533.7 ± 1414.0% in BioFloat, with well-level coefficients of variation of 10.88% and 55.81% respectively; the difference in variance was significant (F-test, F = 50.98, p < 0.0001) (Figure 5B). Whole-plate imaging showed the origin of the difference: signal in NaviPlate was spatially uniform, whereas BioFloat plates displayed weak and heterogeneous signal across wells with no clear positional pattern (Figure 5D,E). Fluorescence microscopy localised BoneTag680 to the spheroid body rather than the well surface or surrounding medium (Figure 5D, inset).

**Figure 5.**
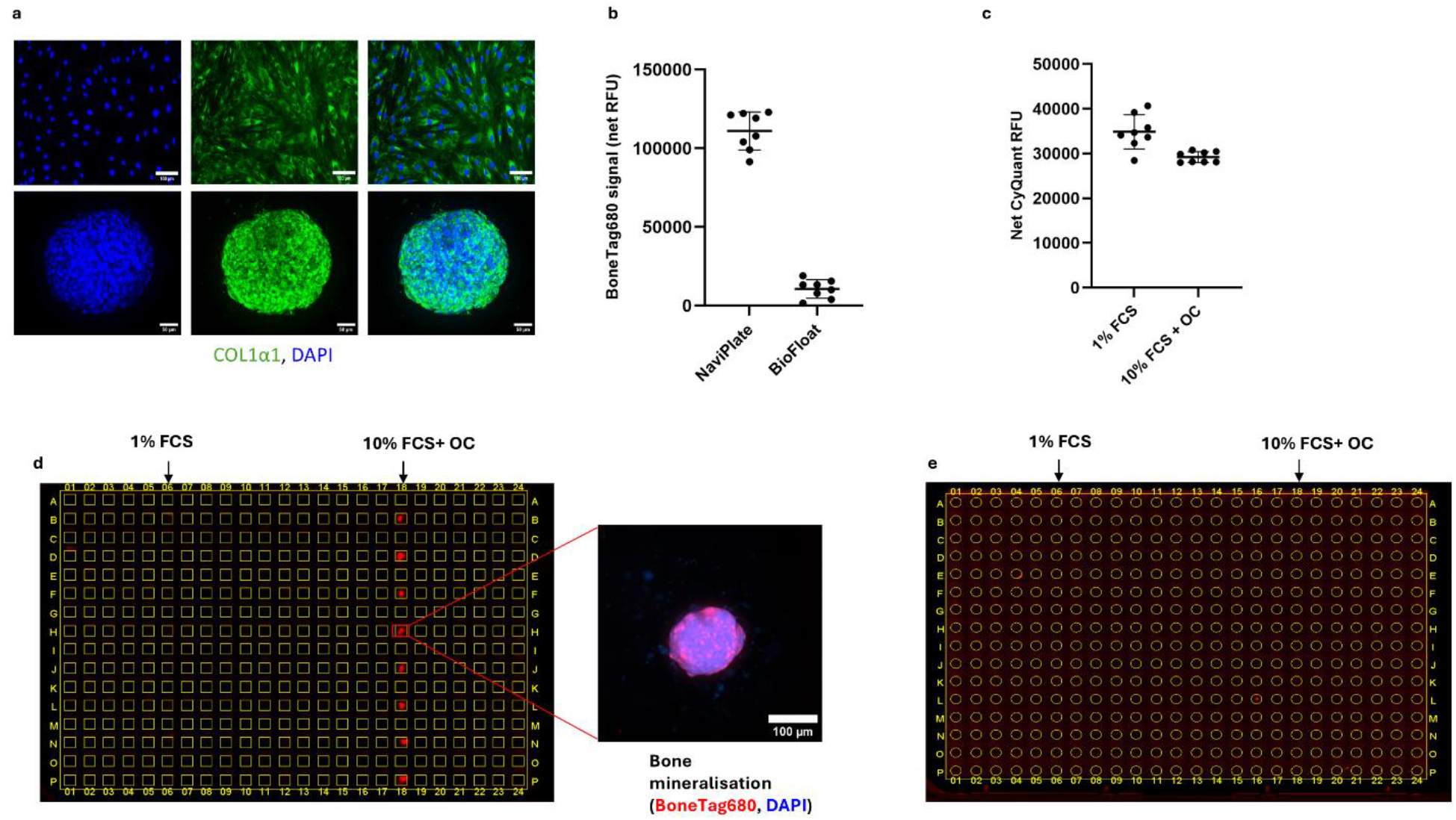
Quantitative mineralisation in DPSC spheroids in the 384-well NaviPlate format. All panels, single donor. (A) COL1α1 immunofluorescence after 24 h in 2D monolayer (top) and 3D spheroid (bottom); DAPI, COL1α1, merge. Scale bars, 100 μm (2D) and 50 μm (3D). (B) BoneTag680 signal in 10% FCS + OC spheroids as a percentage of the 1% FCS condition, NaviPlate versus BioFloat. Points, individual wells; bars, mean ± SD; dashed line, 1% FCS level. n = 8 wells per format; CV = 10.88% and 55.81%; F-test on variances, F = 50.98, p < 0.0001. (C) Net CyQUANT NF fluorescence in 1% FCS and 10% FCS + OC in NaviPlate. (D, E) Whole-plate Odyssey CLx scans, NaviPlate (D) and BioFloat (E); inset in (D), BoneTag680 (red) and DAPI (blue) in a single spheroid. (F) Within-donor assay reproducibility across four independent NaviPlate runs: assay window 13.74 ± 3.29-fold, inter-plate CV 10.24%, Z′ = 0.627 ± 0.119.

To establish that increased signal did not reflect increased biomass, DNA content was measured in the same wells immediately after BoneTag680 acquisition. Net CyQUANT NF fluorescence was 34,851.7 ± 3,836.4 RFU in 1% FCS and 29,196.7 ± 1,219.0 RFU in 10% FCS + OC (Figure 5C). Osteogenic culture therefore produced no increase in cell number - DNA content was approximately 16% lower - while BoneTag680 signal in the same wells was approximately 18-fold higher. Mineral deposition and cell number are thus separable in this format, and both are measurable sequentially from the same well.

Assay reproducibility was assessed across four independent NaviPlate runs performed on separate days from independently thawed stocks. The assay window was 13.74 ± 3.29-fold, with an inter-plate coefficient of variation of 10.24% and a Z′ factor of 0.627 ± 0.119 (Figure 5F). These values describe technical and within-donor reproducibility; donor-to-donor reproducibility was not assessed.

#### Osteogenic marker proteins are detectable within intact spheroids

Spheroids were examined for osteogenic marker expression at days 5 and 7 (Figure 6A-H). COL1α1 was detected in both conditions at both timepoints (Figure 6A,B,E,F), confirming that matrix collagen is retained within intact aggregates and remains accessible to antibody under the in-plate staining protocol. Osteocalcin was likewise detected in both conditions, with heterogeneous distribution between aggregates (Figure 6C,D,G,H). Neither marker discriminated the two conditions at these timepoints under the imaging conditions used. These data are therefore reported as confirmation that marker proteins are detectable within intact spheroids using the in-plate protocol - a methodological result relevant to anyone adapting immunofluorescence to this format - rather than as independent evidence of induction, for which BoneTag680 is the primary measure.

**Figure 6.**
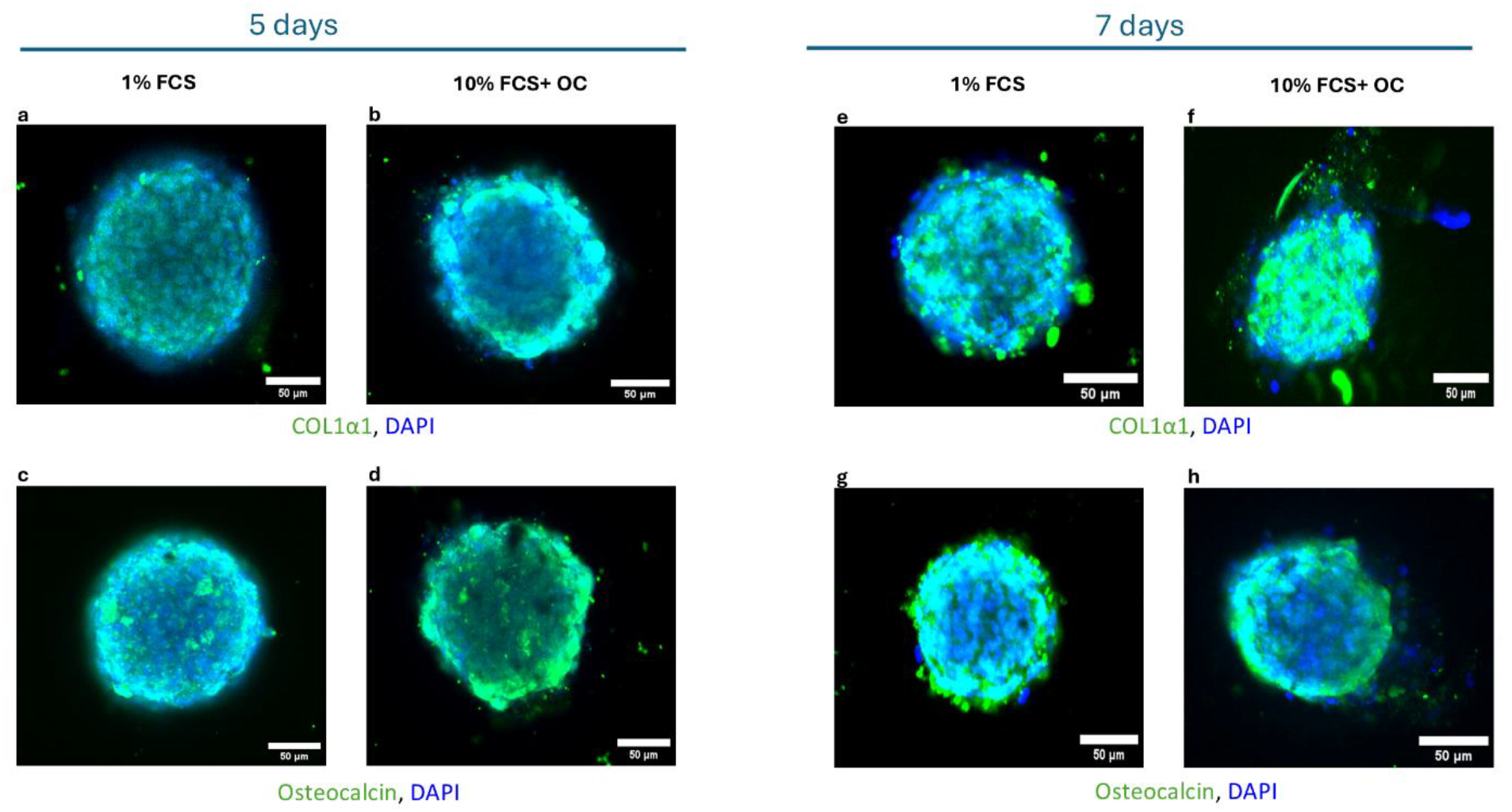
Immunofluorescence analysis of osteogenic markers in DPSC spheroids. (A–D) Day 5: COL1α1 (green; A, 1% FCS; B, 10% FCS + OC) and osteocalcin (green; C, 1% FCS; D, 10% FCS + OC), with DAPI (blue). (E–H) Day 7: COL1α1 (E, F) and osteocalcin (G, H). Representative images; no intensity quantification was performed. Single donor. Scale bars, 100 μm.

#### The format tolerates mechanical perturbation

Plates were placed on an orbital shaker for 4 h per day and spheroids stained for osterix and sclerostin at day 5. Spheroids remained intact and centrally positioned throughout, confirming that the format tolerates orbital agitation without disaggregation or displacement. Osterix- and sclerostin-associated signal appeared more prominent under rocking conditions (Supplementary Figure S2). These observations are qualitative, acquired without intensity quantification, and the hydrodynamic environment was not characterised: orbital agitation generates a spatially non-uniform flow field rather than a defined shear stress. They are reported as evidence that the format is compatible with mechanical stimulation.

## Discussion

We set out to convert 3D human mineralisation from a terminally stained observation into a continuous, well-level quantity. The assay enables primary human DPSCs to form a single, spatially defined spheroid in each well of a 384-well plate; mineral deposition occurs within 5 days and is measured non-destructively by direct in-well near-infrared labelling and plate-reader acquisition. The same plate supports whole-plate imaging and orthogonal DNA-content measurement. The contribution is the integration - spheroid consistency, automated handling and a quantitative optical endpoint made mutually compatible at 384-well density - rather than the demonstration of DPSC osteogenic competence, which is established.^4,5,9^

Two design decisions carried most of the assay performance. The first was that no step involve spheroid transfer. Aggregate handling is a principal source of positional and morphological variance in 3D multiwell workflows,^12^ and eliminating it removes that variance from the measurement rather than correcting for it downstream. The second was the choice of endpoint. Alizarin Red S remains the field standard but requires fixation and extraction for quantification, foreclosing longitudinal measurement. Adding a tetracycline-derived near-infrared probe^13,14^ directly to the culture volume and reading from below converts that principle into a plate-scale measurement, while the near-infrared channel avoids the autofluorescence that limits visible-wavelength detection in dense aggregates.

The objective in defining the two culture conditions was not to partition the contribution of individual medium components, but to establish a pair of anchors. The low anchor required minimal mineral signal with spheroid integrity and cell number maintained across the assay period, which 1% FCS without osteogenic cocktail achieved. The high anchor required maximal mineral signal without an accompanying increase in cell number. Serum concentration proved a determinant of the achievable maximum: spheroids cultured in 1% FCS with osteogenic cocktail produced only a small fraction of the signal obtained in 10% FCS + OC (Supplementary Figure S1), so the cocktail alone does not drive mineralisation at reduced serum. The two conditions should be read as the empirical limits of the assay’s signal range rather than as a matched comparison attributing mineralisation to any one component; formally partitioning the effects of serum and cocktail would require a factorial design and was not attempted.

Because mineral signal and DNA content are acquired from the same well, the assay reports two independent compound effects in a single run. A genuine anabolic candidate should raise mineral signal without raising cell number, as the osteogenic cocktail does here. A compound raising both is more plausibly acting on cell number than on matrix mineralisation, and one reducing DNA content is flagged as cytotoxic at that concentration. The orthogonal channel therefore functions as a built-in counter-screen against a common source of false positives in mineralisation assays, where signal scales with the amount of cell material present. Two caveats bound this: CyQUANT NF reports DNA content at a single timepoint, a proxy for cell number rather than a direct measure of proliferation; and the sensitivity of the channel has not been established against reference proliferative or cytotoxic compounds.

Spheroid formation was strongly format-dependent, consistent with reports that aggregation method, surface chemistry and well geometry determine aggregate size and morphology.^11^ The platform comparison was a qualitative screen without prospectively defined acceptance criteria, and should be read as workflow optimisation under one set of conditions rather than a general ranking of plate products; seeding density, dispense method and cell source all interact with format. The transferable observation is that surface coating rather than geometry alone determined the outcome. Format selection ultimately rested on the quantitative comparison of signal variance.

Several limitations constrain interpretation, the most consequential concerning generality. All data derive from a single DPSC donor; reproducibility is established across four independent runs and describes technical and within-donor performance. Donor-to-donor variability, the dominant source of variance in primary human stromal cell assays, has not been assessed, and establishing the assay window across independent donors is a prerequisite for screening application. Probe specificity also requires direct demonstration: loss of signal under pharmacological inhibition of mineralisation would establish that BoneTag680 reports mineral rather than matrix density or probe retention. The platform comparison was qualitative; immunofluorescence data are qualitative and constrained by antibody penetration and optical depth in intact aggregates; and the mechanical-stimulation observations demonstrate format compatibility without characterising the hydrodynamic environment.

The system should be understood as a defined quantitative assay rather than a screening-validated platform. Screening deployment requires the assay window to be established across independent donors, predefined acceptance criteria, and confirmation that a reference anabolic compound produces the expected concentration-dependent response. It also requires a decision about which condition compounds are tested in: the anchors defined here bracket the signal range but are its extremes, and detecting compounds that enhance mineralisation will require a sub-maximal condition positioned between them.

With those additions, the non-destructive endpoint and 384-well density make the approach directly applicable to compound screening for bone-anabolic activity in a primary human system, in a field where mineralisation is still assessed largely by destructive terminal assays. Given the accessibility of dental pulp as a cell source and the compatibility of the format with cryopreserved banked cells, extension across donor panels is technically straightforward and is the natural next step.

## Supplementary figure legends

**Supplementary Figure S1.**
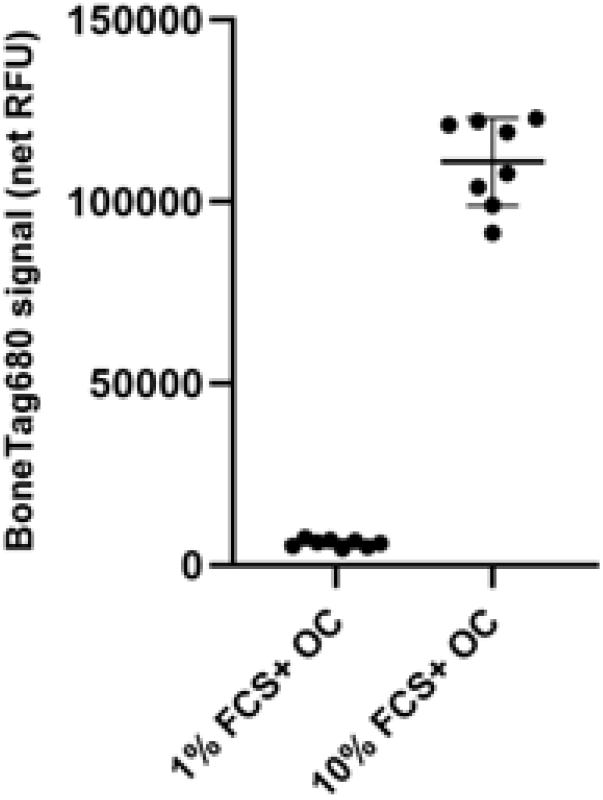
Serum concentration determines the maximal mineralisation achievable with the osteogenic cocktail. Optimisation of the maximal-mineralisation anchor condition. NaviPlate spheroids were maintained for 5 days in 1% FCS + OC or 10% FCS + OC and labelled in-well with BoneTag680. Net BoneTag680 signal was 6,055.9 ± 1,035.9 RFU in 1% FCS + OC and 111,008.3 ± 12,081.4 RFU in 10% FCS + OC. Points represent individual wells; bars show mean ± SD. n = 8 wells per condition, single donor, 3 independent experiments. The osteogenic cocktail alone therefore does not generate mineral signal at reduced serum concentration, and 10% FCS + OC was adopted as the maximal-mineralisation anchor condition.

**Supplementary Figure S2.**
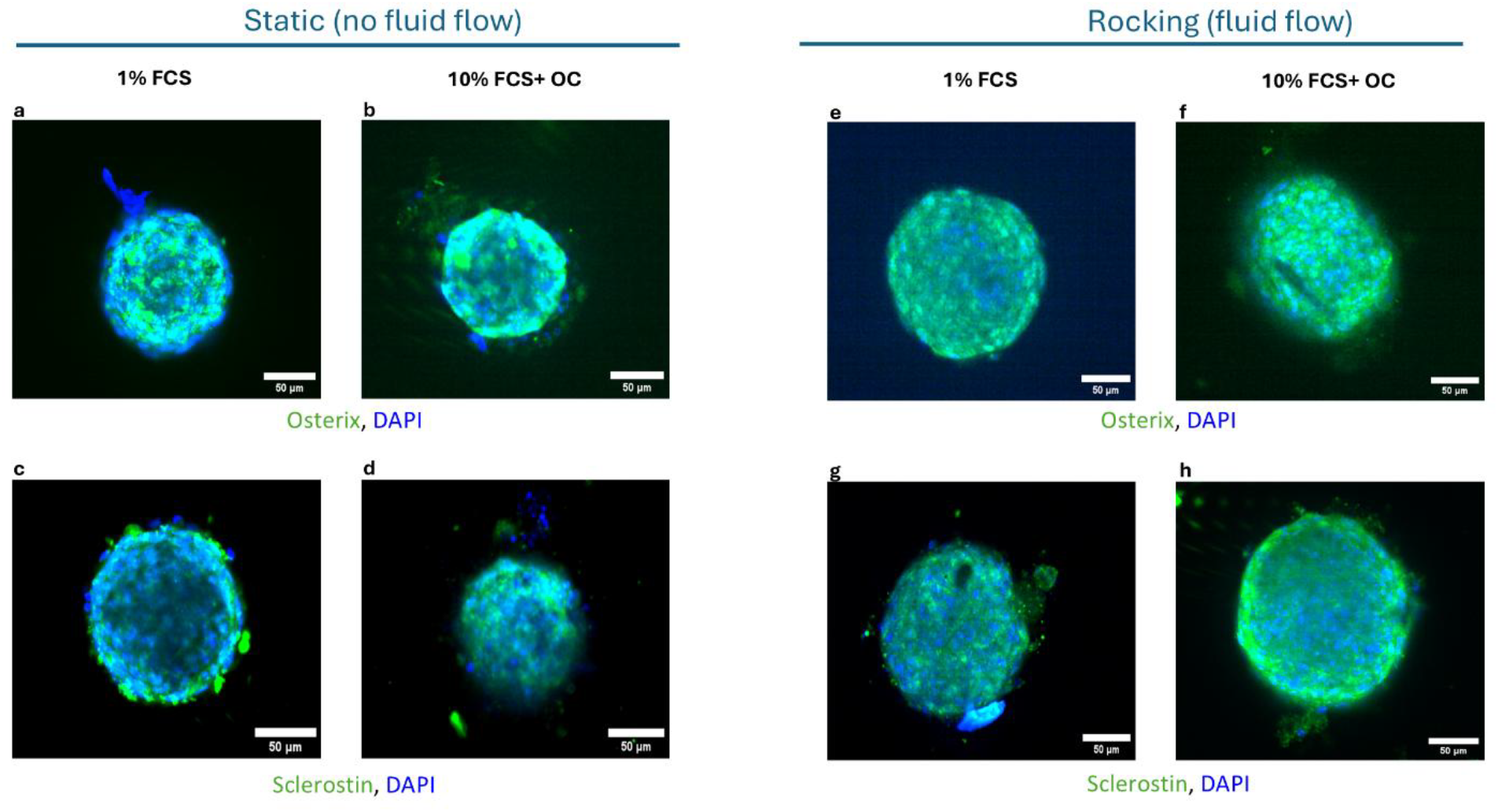
Pilot analysis of osteocyte-associated markers under static and orbital agitation conditions. (A–D) Static: osterix (green; A, 1% FCS; B, 10% FCS + OC) and sclerostin (green; C, 1% FCS; D, 10% FCS + OC), DAPI (blue). (E–H) Agitated: osterix (E, F) and sclerostin (G, H). Representative images from 8 spheroids; no quantitative analysis was performed. Scale bars, 50 µm.

## Acknowledgements

The authors thank Jacqui Cox, Claire Cunningham and Karen Howat for technical support, and acknowledge the MRC Protein Phosphorylation and Ubiquitylation Unit, the High-Content Imaging Facility and the Flow Cytometry Facility at the University of Dundee.

## Author contributions

**Mohammed Alshadidi:** Investigation; Formal analysis; Writing - original draft; Funding acquisition. **Elad Katz:** Methodology; Formal analysis; Supervision; Writing - review and editing. **Ian Ellis:** Supervision; Writing - review and editing. **Sarah Jones:** Supervision; Writing - review and editing. **Mohammad Islam:** Conceptualization; Formal analysis; Supervision; Writing - review and editing.

## Statements and declarations

### Ethical considerations

Human dental pulp tissue was collected with informed consent under Tayside Biorepository delegated ethics (Research Tissue Collection, approval no. TR000620), from third molars extracted during routine clinical procedures that would otherwise have been discarded.

### Consent to participate

Informed consent was obtained from all tissue donors under the approval referenced above.

### Consent for publication

Not applicable. No identifiable individual data are reported.

### Declaration of conflicting interest

E.K. is the inventor of the NaviPlate used in this work and is co-founder and Chief Technology Officer of Navigate Precision Biology Limited, which commercialises the NaviPlate-based assay. M.I. is co-founder and Chief Scientific Officer of Navigate Precision Biology Limited. The University of Dundee holds equity in Navigate Precision Biology Limited. M.A., I.E. and S.J. declare no competing interests. The plate comparison reported in Figure 4 was a qualitative screen; selection of the format used for subsequent assay development was based on the quantitative signal-variance data reported in Figure 5.

### Funding statement

This work was funded by a Government of the Kingdom of Saudi Arabia PhD Scholarship (D-2324862) awarded to M.A.

### Data availability

Raw plate-reader measurements and imaging data are available from the corresponding author on reasonable request. Image analysis used ImageJ v1.52n and quantitative analysis GraphPad Prism v10; plate-reader data were acquired and exported using the manufacturer’s PHERAstar FS software. No custom code was used.

### Materials availability

The NaviPlate 384-well plate is a commercially available product of Navigate Precision Biology Limited and is available to the research community by prior agreement with the company. This study generated no other new unique reagents.

### Use of generative AI

The authors did not use generative AI or AI-assisted technologies in the writing process.

